# Psychedelic drug action at dendrites is gated by behavioral state and serotonin receptors

**DOI:** 10.64898/2026.07.16.738983

**Authors:** Neil K. Savalia, Ling-Xiao Shao, Cory A. Knox, Amelia D. Gilbert, Quan Jiang, Alex C. Kwan

## Abstract

How psychedelics act on cortical dendrites to produce long-lasting structural plasticity remains poorly understood. Here, we characterize the effects of psilocybin on dendritic calcium dynamics in pyramidal tract neurons of the mouse medial frontal cortex. Psilocybin transiently increases calcium event rates in apical dendritic tufts over a time course that parallels the drug’s pharmacokinetics in the brain. This acute effect is brain state-dependent, occurring selectively during quiet wakefulness, and was abolished by cell type-specific deletion of the 5-HT_2A_ receptor. Under control conditions, dendritic calcium signaling predicts subsequent spine formation, but this relationship is not preserved following psilocybin administration. Together, these findings reveal that psilocybin engages brain state- and 5-HT_2A_ receptor-dependent dendritic signaling, while altering the relationship between acute dendritic activity and long-term structural plasticity. The results suggest that the mechanisms linking acute dendritic signaling to structural remodeling differ between physiological and psychedelic-induced plasticity.

## INTRODUCTION

Psychedelics show promise as therapeutics for various psychiatric conditions [1, 2]. One or two treatments with psilocybin, given with psychological support, have been shown to rapidly reduce symptoms of depression with effects lasting for weeks to months [3–7]. Yet, it remains a mystery how a drug that clears from the body within hours prompts such durable effects on mood and behavior. A leading hypothesis is that psychedelics promote long-lasting structural plasticity by driving synaptogenesis in cortical pyramidal neurons [8–10]. Indeed, numerous studies have shown that a single dose of psilocybin can lead to increased density of dendritic spines in the medial frontal cortex [11–13]. Because most excitatory synapses form on dendrites, this process is presumably initiated by the acute effects of psychedelics on dendrites [14]. However, little is known about how psychedelics alter dendritic physiology immediately after administration or how these acute effects relate to subsequent structural remodeling. Delineating the mechanistic link may provide insights into whether the acute effects of psychedelics are necessary for their long-term therapeutic benefits [15–17].

Decades of research have identified dendritic calcium signaling as a central mechanism underlying synaptic plasticity. Calcium acts as a second messenger that links neuronal activity to the molecular pathways governing synapse formation and remodeling [18, 19]. In pyramidal neurons, depolarization triggers calcium influx into dendrites. Postsynaptic calcium has been shown to be necessary, sufficient, and physiologically relevant for the induction of synaptic plasticity [20, 21]. Consistent with this framework, in vivo imaging studies have demonstrated that dendritic calcium signaling precedes plasticity in cortical pyramidal neurons, which may involve both branch-specific calcium transients [22] and more global dendritic plateau potentials [23]. These findings support acute changes in dendritic calcium signaling as a critical early step towards structural plasticity, although whether the relationship holds for psychedelics is unclear.

We expect serotonin receptor signaling and behavioral state to be key determinants for psilocybin’s acute actions on dendrites. Psilocybin is rapidly metabolized to psilocin, which acts as an agonist at multiple serotonin receptor subtypes [24]. The 5-HT_2A_ receptor is required for the acute subjective effects of psychedelics [25] and is highly expressed in the apical dendrites of cortical pyramidal neurons [26, 27], whereas the 5-HT_1A_ receptor is also expressed in these neurons and modulates the acute behavioral effects of psilocybin [28]. Because these receptors exert opposing effects on neuronal excitability [29, 30], they may act in concert to shape dendritic responses to the drug. In parallel, the acute actions of psychedelics are thought to depend strongly on internal state and environmental context, often referred to as “set and setting” [31–35]. Consistent with this idea, recent studies have shown that psychedelic-evoked neural responses are enhanced during low-demand behavioral states in rodents [36] and humans [37]. Apical tuft dendrites are thought to integrate contextual information and contribute to conscious perception [38–42], making them a plausible neural substrate for the state-dependent effects of psychedelics. We therefore asked how behavioral state influences psilocybin-evoked dendritic activity and whether the effects are mediated by specific serotonin receptor subtypes.

## RESULTS

### Psilocybin induces a transient increase in spontaneous dendritic calcium events in frontal cortical PT neurons

In this study, we focused on layer 5 pyramidal tract (PT) subtype of pyramidal cells in the mouse medial frontal cortex, which have elaborated apical dendritic arborization, and were shown to exhibit psilocybin-induced structural neural plasticity and contribute to the drug’s long-term behavioral effects [13]. PT neurons are primarily subcerebral projection neurons that send axons to subcortical regions such as the thalamus, midbrain, and pons [43, 44]. To target the frontal cortical PT neurons, adult C57BL/6J mice were injected with AAV1-CAG-Flex-mRuby2-GSG-P2A-GCaMP6f-WPRE-pA in the medial frontal cortex and a low titer of AAVretro-hSyn-Cre-WPRE-hGH in the pons (**Figure 1A**). This approach enabled sparse labeling of a small number of PT neurons with a static red fluorophore mRuby2 for structural imaging and a genetically encoded calcium indicator GCaMP6f for calcium imaging [45] (**Figure 1B**). Mice were implanted with a chronic glass window to allow for longitudinal imaging in head-fixed mice habituated to run on a linear treadmill.

**Figure 1.**
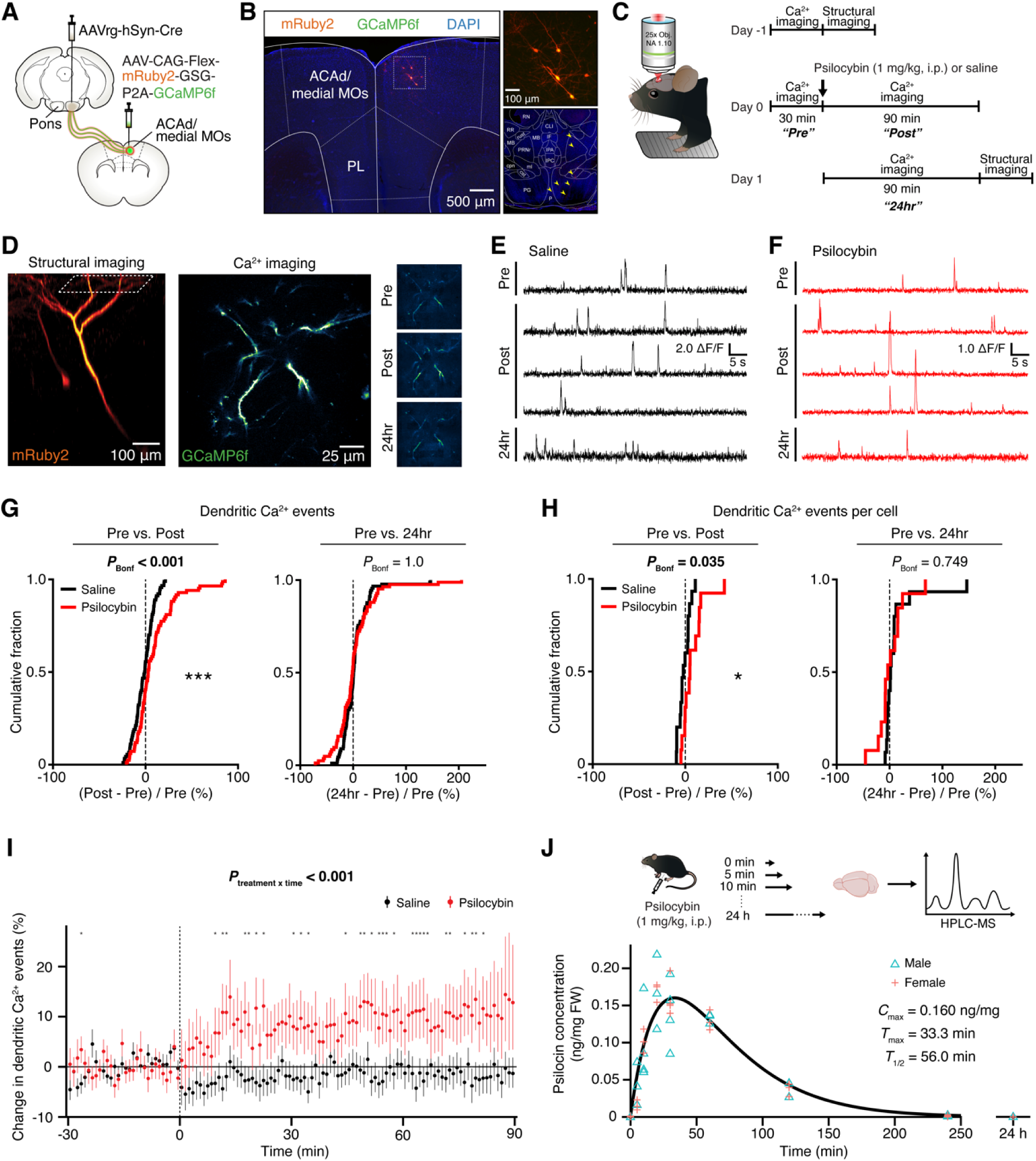
Psilocybin transiently increases the spontaneous rate of dendritic calcium events in frontal cortical PT pyramidal neurons. **(A)** Schematic of the strategy to co-express mRuby2 and GCaMP6f in PT neurons in the medial frontal cortex of C57BL/6J mice. **(B)** Histological images of sparsely labeled PT neurons in medial frontal cortex. Insets, cell bodies in the medial frontal cortex and axons in the pons. **(C)** Timeline of the two-photon microscopy experiment. **(D)** In vivo two-photon imaging, including dendritic arborization from z-stack images and field of view containing GCaMP6f-expressing dendrites. **(E)** Time-lapse Δ*F*/*F*_0_ from a dendritic branch before and after saline treatment. **(F)** Similar to E, but for psilocybin (1 mg/kg, i.p.). **(G)** Cumulative distribution plot of the fractional change in the rate of calcium events detected in PT dendritic branches after psilocybin (red) or saline (black) acutely after treatment (“Post”) or a day later (“24hr”) relative to pre-treatment baseline (“Pre”). *P* values from post-hoc *t*-tests were adjusted using Bonferroni correction for multiple comparisons. ***, *P* <□0.001. **(H)** Similar to G, but with dendritic branch data averaged per neuron. *, *P* <□0.05. **(I)** Minute-by-minute fractional change in the rate of calcium events detected in PT dendritic branches for psilocybin (red) or saline (black) on the day of treatment. Circle, group mean. Error bars, the bootstrapped 95% confidence interval of the mean. Asterisk denotes *P* < 0.05 on *post hoc* two-sample *t*-test (psilocybin vs. saline) after Bonferroni correction for multiple comparisons across time. **(J)** Brain psilocin concentration measured using high-performance liquid chromatography-tandem mass spectrometry (HPLC-MS) after treatment with psilocybin (1 mg/kg, i.p.). A one-compartment model was used to estimate time to peak (*T*_max_), peak concentration (*C*_max_), and elimination half-life (*T*_1/2_). For (G)–(I), n = 84 and 90 dendritic branches from 13 and 15 PT neurons for psilocybin and saline, respectively. For (J), n = 6-8 mice per time point. Full statistical analyses are provided in **Table S1**.

The apical tuft dendrites of frontal cortical PT neurons were imaged for three days centered on a single treatment with psilocybin (1 mg/kg, i.p.) or an equivalent volume of saline (**Figure 1C**). On day -1, baseline imaging was used to locate a field of view suitable for dendritic calcium imaging and to acquire a z-stack of the dendritic structure. On day 0, the same field of view was imaged continuously at 30 Hz for 30 minutes before (“Pre”) then 90 minutes immediately after treatment (“Post”). On day 1, an additional 60-90 minutes of calcium imaging (“24hr”) were collected in the same field of view, before acquiring another z-stack of the dendritic structure.

For structural imaging, mice were lightly anesthetized to reduce motion artifacts. Because of the sparse labeling, most fields of view contained dendrites from a single cell (**Figure 1D**). The highly restricted expression enabled definitive assignment of each dendrite to its parent neuron and reliable tracking of the same dendrites across imaging sessions (**Figure 1E, F**).

We imaged apical tuft dendrites from 28 PT neurons (13 for psilocybin and 15 for saline) in 21 mice (11 females and 10 males). In total, we tracked 174 dendritic branches (84 for psilocybin and 90 for saline). Calcium events were inferred from the fluorescence signals using a machine-learning algorithm [46]. To account for the hierarchical nature of the dataset, all statistical tests were performed using linear mixed-effects models including a random-effects term for dendrites nested within cells and mice (see **Table S1**). The rate of spontaneous dendritic calcium events increased acutely following psilocybin administration, but returned to baseline levels by 24 hours (treatment × time interaction: *P* = 0.006, linear mixed-effects model with dendrites nested within cells and mice). Compared with baseline, psilocybin transiently increased the rate of dendritic calcium events by approximately 10% (psilocybin: 8.9±2.4%, mean±SEM; saline: -1.8±1.1%; Bonferroni-corrected post hoc comparison: *P* = 9 x 10^-5^; **Figure 1G**). This effect was absent 24 hours later (psilocybin: 2.6±4.3%; saline: 4.5±2.9%). Averaging across dendritic branches within each neuron yielded qualitatively similar results, indicating that the acute increase was also evident at the level of individual cells (**Figure 1H**).

### The dendritic calcium dynamics track the brain pharmacokinetics of psilocybin

As a function of time, the psilocybin-evoked increase in dendritic calcium events developed over the first 10 minutes and remained elevated for the remainder of the imaging session (treatment × time interaction: *P* = 2 x 10^-6^; **Figure 1I**). To determine whether this time course reflected the pharmacokinetics of the drug, we quantified brain psilocin concentrations in adult C57BL/6J mice using tandem high-performance liquid chromatography and mass spectrometry. Psilocin levels were measured at 5, 10, 20, 30 min, and 1, 2, 4, and 24 h after psilocybin administration (1 mg/kg, i.p.), together with untreated controls (n = 60 mice including 29 females and 31 males; 6-8 mice per time point). As expected, brain psilocin levels rose quickly following psilocybin administration, before declining to undetectable levels by 24 hours (**Figure 1J**). Fitting the data with a one-compartment pharmacokinetics model yielded a time to peak concentration of 33 min and an elimination half-life of 56 min. The peak brain concentration was 0.16 ng/mg, corresponding to approximately 0.81 μM. Notably, the time course of the increase in dendritic calcium events closely tracked brain psilocin concentrations, with a significant positive correlation (*P* = 0.02; **Figure S1**). By contrast, comparison with the time course of the head-twitch response from our previous study [47] showed that both brain psilocin concentrations and dendritic calcium activity lagged behind the behavioral response, which peaks 6-8 minutes after psilocybin administration (**Figure S1**). Together, these results show that psilocybin produces a transient increase in spontaneous dendritic calcium activity whose temporal profile closely mirrors drug exposure in the brain.

### Acute dendritic calcium responses to psilocybin require 5-HT_2A_ receptors

Next, we investigated the role of distinct 5-HT receptor subtypes in mediating the psilocybin-evoked increase in dendritic calcium activity. Frontal cortical PT neurons express both 5-HT_2A_ and 5-HT_1A_ receptors [13], which are main binding targets of psilocin [24]. For PT neuron-selective receptor knockout of 5-HT_2A_ receptor, we used a *Htr2a^f/f^* mouse [13, 48], injecting AAV1-CAG-Flex-mRuby2-GSG-P2A-GCaMP6f-WPRE-pA in the medial frontal cortex and a low titer of AAVretro-hSyn-Cre-WPRE-hGH in the pons (**Figure 2A**). We imaged apical tuft dendrites from 18 PT neurons (11 for psilocybin and 7 for saline) in 9 *Htr2a^f/f^*mice (7 females and 2 males), tracking 132 dendritic branches (87 for psilocybin and 45 for saline) (**Figure 2B**). In PT neurons without 5-HT_2A_ receptors, psilocybin (1 mg/kg, i.p.) no longer altered the spontaneous rate of dendritic calcium events acutely after treatment (psilocybin: 3.1±1.6%; saline: 0.3±1.6%; Bonferroni-corrected post hoc comparison: *P* = 0.5; **Figure 2C**). Intriguingly, psilocybin-treated mice with the cell type-specific 5-HT_2A_ receptor knockout showed reduced dendritic calcium signaling a day later (psilocybin: -6.6±1.8%; saline: 1.9±2.5%; Bonferroni-corrected post hoc comparison: *P* = 0.014). Overall, because of the next-day suppression of dendritic calcium events, there was a significant treatment × time interaction (*P* = 2 x 10^-4^, linear mixed-effects model with dendrites nested within cells and mice).

**Figure 2.**
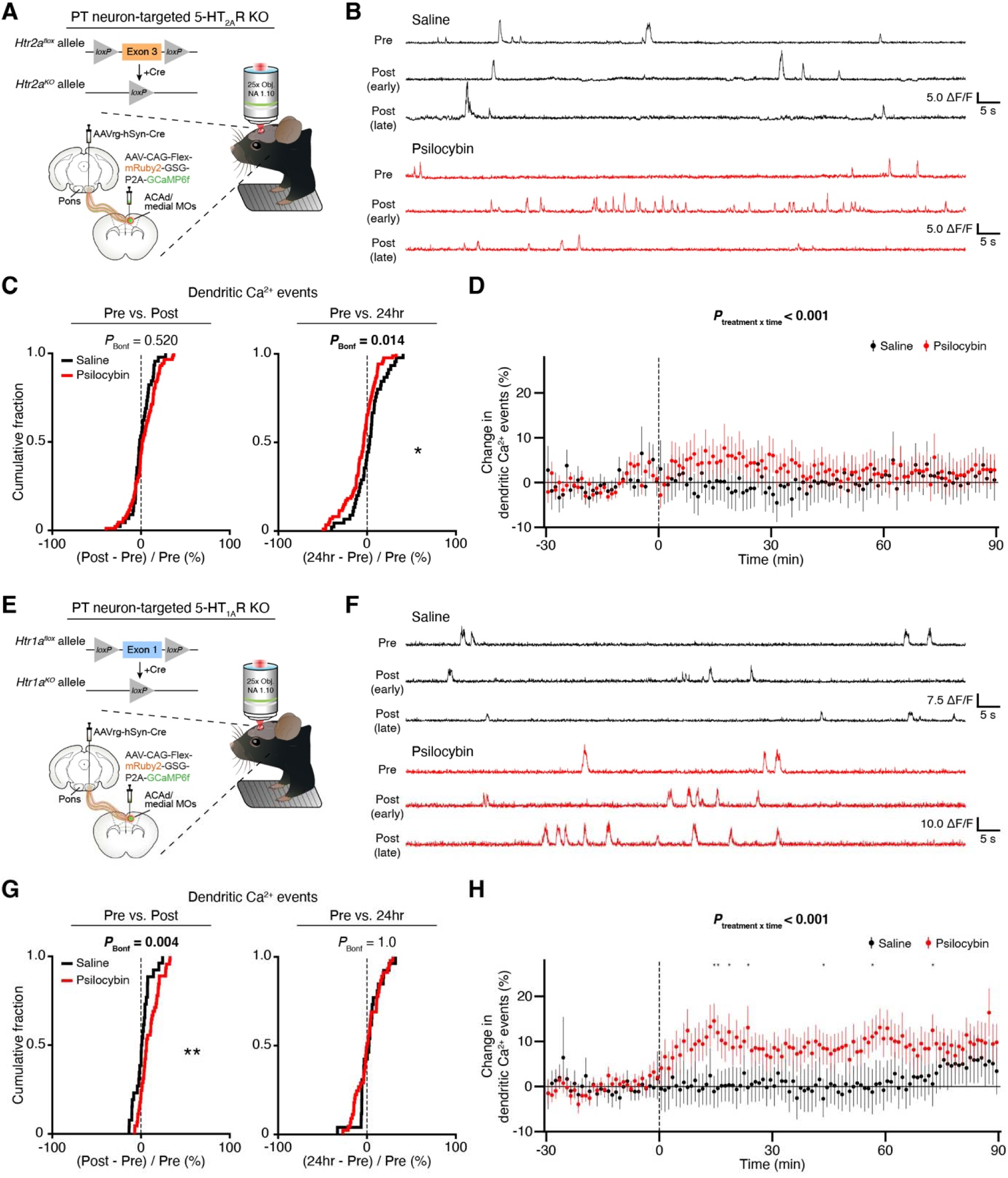
Knockout of 5-HT_2A_, but not 5-HT_1A_, receptors abolishes the psilocybin-evoked increase in dendritic Ca^2+^ dynamics. **(A)** Schematic of the strategy to co-express mRuby2 and GCaMP6f in PT neurons in the medial frontal cortex of *Htr2a^f/f^*mice, while inducing cell type-specific knockout of the 5-HT_2A_ receptor. **(B)** Time-lapse Δ*F*/*F*_0_ from a dendritic branch before and after treatment with saline (black), and another dendritic branch before and after treatment with psilocybin (1 mg/kg, i.p.; red). **(C)** Cumulative distribution plot of the fractional change in the rate of calcium events detected in PT dendritic branches with 5-HT_2A_ receptor knockout after psilocybin (red) or saline (black) acutely after treatment (“Post”) or a day later (“24hr”) relative to pre-treatment baseline (“Pre”). *P* values from post-hoc *t*-tests were adjusted using Bonferroni correction for multiple comparisons. *, *P* < 0.05. **(D)** Minute-by-minute fractional change in the rate of calcium events detected in PT dendritic branches with 5-HT_2A_ receptor knockout for psilocybin (red) or saline (black) on the day of treatment. Circle, group mean. Error bars, the bootstrapped 95% confidence interval of the mean. Asterisk denotes *P* < 0.05 on *post hoc* two-sample *t*-test (psilocybin vs. saline) after Bonferroni correction for multiple comparisons across time. **(E-H)** Similar to A-D, but for PT neurons in the medial frontal cortex of *Htr1a^f/f^* mice, which induced cell type-specific knockout of the 5-HT_1A_ receptor. **, *P* <□0.01. For (A)–(D), n = 87 and 45 dendritic branches from 11 and 7 PT neurons for psilocybin and saline, respectively. For (E)–(H), n = 45 and 26 dendritic branches from 5 and 4 PT neurons for psilocybin and saline, respectively. Full statistical analyses are provided in **Table S1**.

For conditional deletion of the 5-HT_1A_ receptor, we used a *Htr1a^f/f^* mouse [49, 50], again injecting AAV1-CAG-Flex-mRuby2-GSG-P2A-GCaMP6f-WPRE-pA in the medial frontal cortex and a low titer of AAVretro-hSyn-Cre-WPRE-hGH in the pons (**Figure 2E**). We imaged apical tuft dendrites from 9 PT neurons (5 for psilocybin and 4 for saline) in 6 *Htr1a^f/f^*mice (3 females and 3 males), tracking 71 dendritic branches (45 for psilocybin and 26 for saline) (**Figure 2F**). Here, the absence of 5-HT_1A_ receptor did not affect the acute psilocybin-induced increase in dendritic calcium signaling (psilocybin: 9.3±1.6%; saline: 1.2±1.8%; treatment × time interaction: *P* = 0.011, Bonferroni-corrected post hoc comparison: *P* = 0.004, linear mixed-effects model; **Figure 2G**). Psilocybin had no effect on the dendritic calcium dynamics 24 hours later (psilocybin: 0.9±2.0%; saline: 2.6±2.5%). At the minute-by-minute level, psilocybin’s acute effect on dendritic calcium events in *Htr1a^f/^*^f^ mice varied significantly over time (treatment × time interaction: *P* < 0.001, linear mixed-effects model with dendrites nested within cells and mice; **Figure 2H**). Collectively, deletion of 5-HT_2A_ receptor in the specific sparse set of imaged frontal cortical PT neurons abolished the psilocybin-evoked increase in dendritic calcium events. By contrast, the acute effect of psilocybin on dendritic calcium signaling was unchanged after the deletion of 5- HT_1A_ receptor. A full statistical test across genotypes supports this conclusion, showing a a significant three-way interaction of treatment × time × genotype (*P* < 0.001, linear mixed-effects model; **Figure S2**). The data demonstrate that psilocybin’s acute effects on dendritic calcium dynamics depend critically on the 5-HT_2A_ receptor.

**Psilocybin enhances dendritic calcium activity selectively during quiet wakefulness** Internal states and environmental context have long been recognized to influence psychedelic drug action [31–35]. During imaging, head-fixed mice were free to run on a linear treadmill, giving us the measurement of voluntary locomotion to serve as a proxy for behavioral state [51] (**Figure 3A**). Psilocybin reduced locomotor activity in head-fixed C57BL/6J mice. Relative to the pre-drug baseline, distance traveled per 10-minute interval changed by -158±50 m following psilocybin, compared with a change of -64±46 m following saline (mean±SEM; n = 21 mice total; treatment: *P* < 0.001; **Figure S3A**). Interestingly, this locomotor suppression was not evident in *Htr2a^f/f^* and *Htr1a^f/f^*mice (**Figure S3A**). Despite these differences, all animals exhibited alternating periods of running and quiet wakefulness, allowing us to examine psilocybin’s effects on dendritic calcium dynamics across distinct behavioral states.

**Figure 3.**
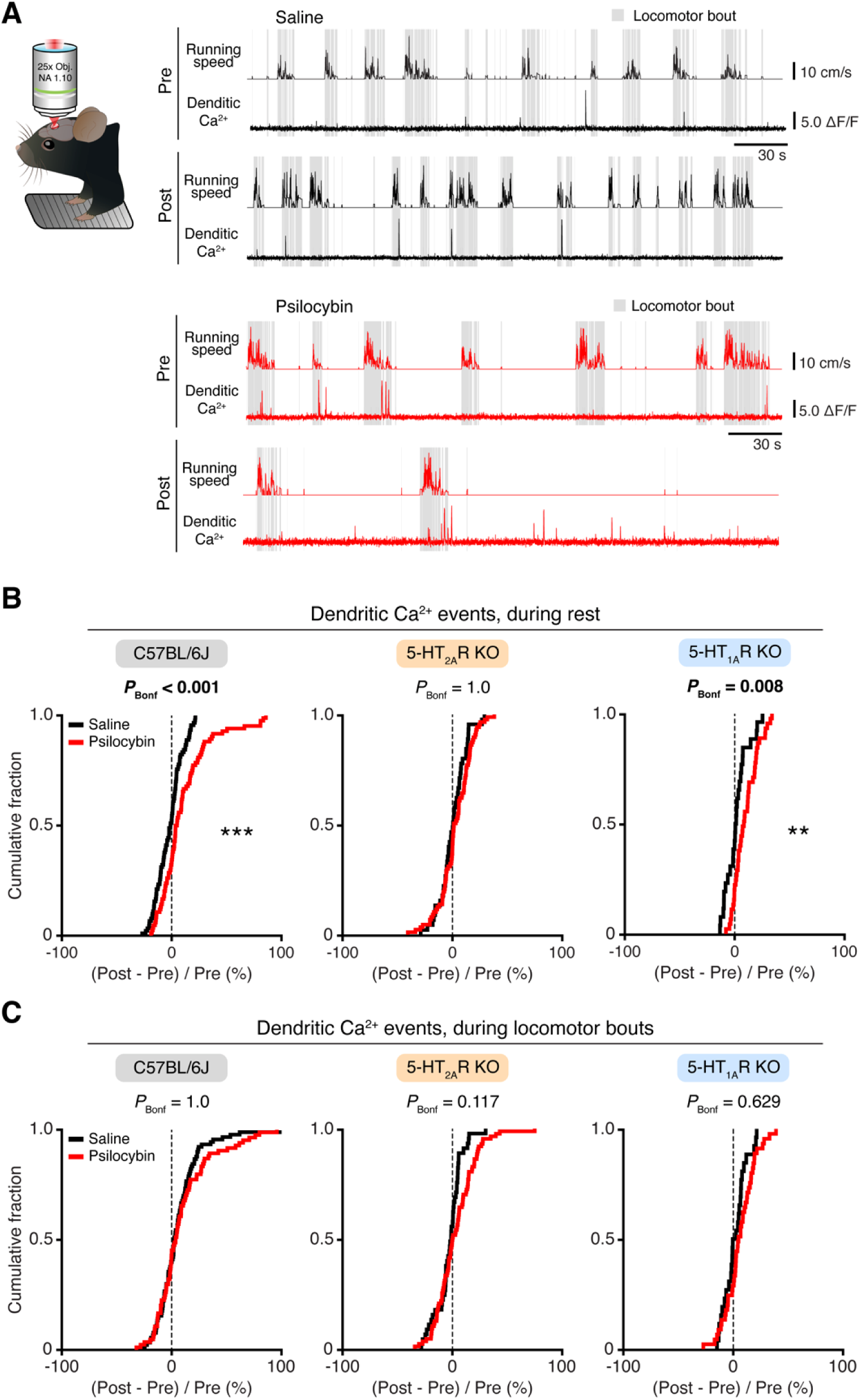
Psilocybin enhances dendritic calcium activity selectively during quiet wakefulness. **(A)** Locomotor activity was monitored with a linear treadmill during imaging sessions. Example pre- and post-treatment locomotor activity and time-lapse Δ*F*/*F*_0_ for a PT dendritic branch in a mouse treated with saline (black) or another PT dendritic branch in a mouse treated with psilocybin (1 mg/kg, i.p., red). Grey shading, run epochs defined as imaging frames with speed >0 cm/s. **(B)** Cumulative distribution plot of the fractional change in the rate of calcium events detected in PT dendritic branches, restricting analyses to rest epochs, after psilocybin (red) or saline (black) acutely after treatment (“Post”) relative to pre-treatment baseline (“Pre”), for C57BL/6J mice (left), *Htr2a^f/f^* mice with 5-HT_2A_ receptor knockout (middle), and *Htr1a^f/f^* mice with 5-HT_1A_ receptor knockout (right). *P* values from post-hoc *t*-tests were adjusted using Bonferroni correction for multiple comparisons. **, *P* < 0.01. ***, *P* <□0.001. **(C)** Similar to B, but restricting analyses to run epochs. For C57BL/6J, n = 84 and 90 dendritic branches from 13 and 15 PT neurons for psilocybin and saline, respectively. For *Htr2a^f/f^*, n = 87 and 45 dendritic branches from 11 and 7 PT neurons for psilocybin and saline, respectively. For *Htr1a^f/f^*, n = 45 and 26 dendritic branches from 5 and 4 PT neurons for psilocybin and saline, respectively. Full statistical analyses are provided in **Table S1**.

We classified behavioral state based on treadmill speed, defining rest and run epochs as 0 and >0 cm/s, respectively. Following psilocybin administration, dendritic calcium activity increased selectively during quiet wakefulness, with a significant effect during rest (psilocybin: 10.5±2.5%; saline: -1.9±1.2%; treatment × locomotion interaction: *P* = 0.019, linear mixed-effects model with dendrites nested within cells and mice; **Figure 3B**), but not during locomotion (psilocybin: 9.1±2.7%; saline: 5.5±2.0%; **Figure 3C**). This behavioral state dependence could not be explained by higher baseline activity during locomotion, as dendritic calcium event rates did not differ between quiescence and locomotion before drug administration (**Figure S3B**).

Conditional deletion of 5-HT_2A_ receptors from PT neurons eliminated the state-dependent response, with no detectable change in dendritic activity during either rest or run (psilocybin, rest: 2.7±1.6%; saline, rest: 0.3±1.8%; psilocybin, run: 3.5±1.9%; saline, run: -3.0±1.8%). By contrast, deletion of 5-HT_1A_ receptors preserved the preferential effect of psilocybin during rest (psilocybin, rest: 9.9±1.6%; saline, rest: 1.6±2.0%; psilocybin, run: 6.0±2.0%; saline, run: 1.6±2.0%). The dependence on behavioral state was not observed 24 hours after psilocybin administration (**Figure S4**). These findings show that psilocybin engages 5-HT_2A_ receptor-dependent dendritic calcium signaling preferentially during quiet wakefulness.

### Psilocybin alters the relationship between dendritic calcium activity and structural plasticity

A main goal of the study was to relate acute dendritic activity to subsequent structural remodeling within the same dendritic branch. For this, we used a bicistronic construct to co-express the static fluorophore mRuby2 and the genetically encoded calcium indicator GCaMP6f within the same sparse set of PT neurons (**Figure 4A**). The mRuby2 fluorescence provided stable structural labeling for quantifying spine formation and elimination between day -1 to day 1, allowing us to directly relate changes in spine density to dendritic calcium activity measured on day 0 (**Figure 4B**).

**Figure 4.**
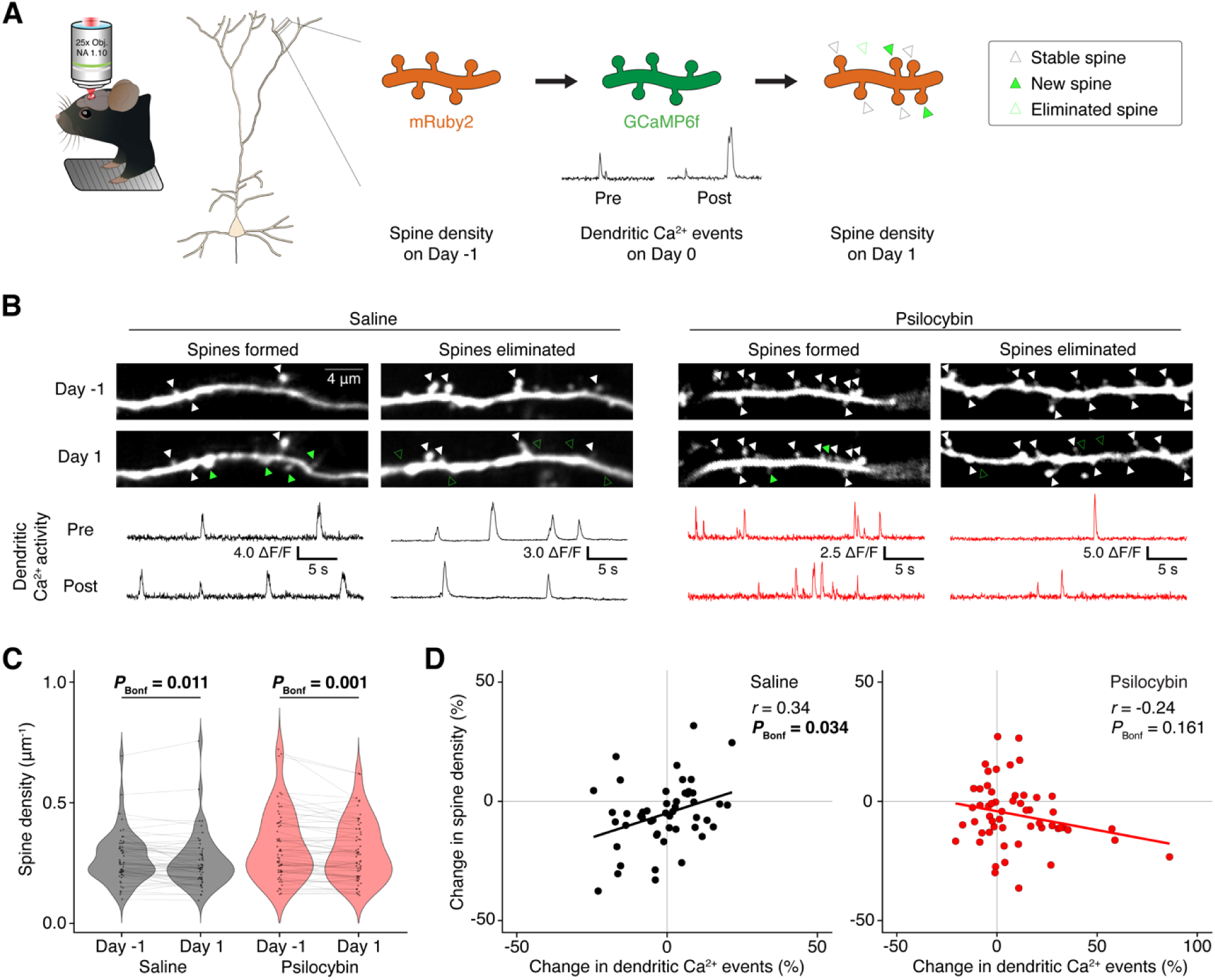
Psilocybin transiently increases the spontaneous rate of dendritic calcium events in frontal cortical PT pyramidal neurons. **(A)** The experimental design allowed for within-dendrite comparison of psilocybin’s acute actions on dendritic calcium activity and subsequent dendritic spine turnover. **(B)** Example dendritic branches with spine formation and elimination in saline and psilocybin groups, and the corresponding dendritic calcium activity in the same dendritic branches on day 0. **(C)** Dendritic spine density on day -1 and day 1 for each dendritic branch in saline and psilocybin groups. **(D)** Change in dendritic calcium events versus change in spine density, for saline (left) and psilocybin (right) groups. Circle, individual dendritic branch. *r*, Pearson correlation coefficient. For (C), n = 60 and 53 dendritic branches from 11 and 10 PT neurons for psilocybin and saline, respectively. For (D), n = 55 and 50 dendritic branches from 11 and 10 PT neurons for psilocybin and saline, respectively. Full statistical analyses are provided in **Table S1.**

We had longitudinal structural imaging data from most cells in the C57BL/6J mice, including 21 PT neurons (11 for psilocybin and 10 for saline) in 21 mice (11 females and 10 males), tracking 113 dendritic branches (60 for psilocybin and 53 for saline). Unexpectedly, both psilocybin- and saline-treated mice exhibited a reduction in spine density over time (main effect of time: *P* < 0.001, mixed-effects model; **Figure 4C**). Spine density decreased by 6.9% after psilocybin administration (day -1: 0.32±0.02 µm^-1^; day 1: 0.30±0.02 µm^-1^; Bonferroni-corrected post hoc comparison: *P* = 0.0013) and by 5.2% after saline injection (day -1: 0.27±0.01 µm^-1^; day 1: 0.26±0.01 µm^-1^, Bonferroni-corrected post hoc comparison: *P* = 0.011). This result contrasts with our previous studies showing psilocybin-induced spine formation [11, 13]. The reason behind the discrepancy is unclear. Compared to previous studies, we note that mRuby2 was expressed from a bicistronic construct, resulting in noticeably dimmer structural labeling that was further diminished by repeated imaging on consecutive days. In addition, animals were head-fixed and underwent imaging during the acute post-psilocybin period, rather than returning to their home cage, raising the possibility that behavioral context during the acute drug period influences the psychedelic-evoked structural plasticity.

For most of the dendritic branches tracked for structure, we had corresponding dendritic calcium imaging data on day 0, including 105 dendritic branches (55 for psilocybin and 50 for saline) from 21 PT neurons in 21 mice. For each dendritic branch, we compared the change in dendritic spine density from day -1 to day 1 against the change in dendritic calcium events on day 0 (**Figure 4D**). Saline-treated mice showed a significant positive association, indicating that elevated dendritic calcium dynamics was associated with subsequent increase in density spines in that dendritic branch (*r* = 0.34, *P* = 0.034). By contrast, we did not detect any relationship between the psilocybin-evoked change in dendritic calcium signals with subsequent structural remodeling (*r* = -0.24, *P* = 0.16). This interpretation is supported by a mixed-effects model to predict the change in dendritic spine density by the change in dendritic calcium events and treatment, which showed a significant interaction effect (*P* = 0.008). Therefore, dendritic calcium activity predicted later spine formation under control conditions, but this coupling was lost after psilocybin administration. The findings suggest that the acute dendritic responses to psilocybin may be largely orthogonal to the mechanisms that govern its longer-term structural remodeling.

## DISCUSSION

In this study, we characterized the mechanisms underlying psilocybin’s effects on dendritic calcium dynamics. A central finding is that physiological spine formation occurred in dendritic branches with more dendritic calcium activity, but this predictive relationship did not hold for psilocybin. Although both the acute dendritic response and subsequent structural remodeling engage apical dendrites and require functional 5-HT_2A_ receptors [13], our results suggest that they diverge downstream of these shared initial events. The findings add to a growing body of evidence that some acute pharmacological responses, including immediate early gene induction and glutamate efflux, do not necessarily predict the long-term therapeutic actions of psychedelic compounds and their analogs [17].

Another key finding is that the effects of psychedelics on neural dynamics depend on the behavioral state of the animal. Behavioral state is one component of the broader concept of ‘set and setting’, in which an individual’s internal state and environmental context powerfully influence the acute subjective effects of psychedelics [31–35]. We show that psilocybin preferentially increases dendritic activity during periods of quiet wakefulness in a 5-HT_2A_ receptor-dependent manner. This result and related recent human findings [37] suggest that psilocybin’s impact is enhanced during quiescent states. This observation is consistent with current clinical practice, where dosing sessions occur in a minimally stimulating environment [52–54]. Our results provide a potential cellular correlate for the state dependence of psychedelic drug action.

This study has several limitations. Contrary to our previous work [11, 13], we did not observe psilocybin-induced dendritic spinogenesis. One possible explanation is the different behavioral context during the acute drug period, as animals in the present study remained head-fixed for imaging rather than returning to their home cage. Future experiments that systematically manipulate behavioral context during psychedelic administration may help determine how acute experience influences subsequent structural plasticity. In addition, dendritic calcium transients can arise from multiple forms of electrical activity, including excitatory synaptic inputs, local NMDA spikes, dendritic plateau potentials, and backpropagating action potentials [55–57]. Given the event frequency and sensitivity of GCaMP6f, we expect that most calcium events detected in this study reflect widespread dendritic electrogenesis associated with burst firing in the soma [58, 59]. Resolving how psychedelics differentially modulate these forms of dendritic activity and how each contributes to long-term structural remodeling will be an important direction for future research.

In sum, psilocybin engages 5-HT_2A_ receptor-dependent dendritic signaling in frontal cortical pyramidal neurons, with preferential effects during quiet wakefulness. The mechanistic separation between acute dendritic activity and long-term structural plasticity suggests that these processes may be partially dissociable, providing a framework for developing next-generation therapeutics.

## METHODS

### Animals

Wild-type C57BL/6J mice (000664) were purchased from Jackson Laboratory and housed in our animal facility. *Htr2a^f/f^* mice were described in a previous study [48] and bred in our animal facility. *Htr1a^f/f^* mice were described in a prior study [49] and bred in our animal facility. Mice were aged 5-8 weeks when surgical procedures including viral injection began. Glass window implant occurred 2 weeks later. Two-photon imaging then took place 3-4 weeks after window implantation. Therefore, mice were aged 10-14 weeks at the time of imaging. Mice were housed in groups with 2-5 mice per cage in a temperature-controlled room, operating under a 12 h-12 h light-dark cycle (08:00 to 20:00 for light), at 70-72 °F ambient temperature, and 30-70% humidity. Food and water were available ad libitum. Male and female mice were randomly assigned to different experimental groups. Animal care and experimental procedures were conducted in accordance with the ethical standards of the National Institutes of Health and were approved by the Institutional Animal Care & Use Committees (IACUC) at Cornell University.

### Viruses

AAV1-CAG-Flex-mRuby2-GSG-P2A-GCaMP6f-WPRE-pA (68719) and AAVretro-hSyn-Cre-WPRE-hGH (105553) were bought from Addgene. AAVretro is an adeno-associated virus (AAV) designed for efficient retrograde transport [60]. Viruses had titers of ≥2[×[10^13^ viral genomes per mL and were stored at -80 °C. Before stereotaxic injection, viral aliquots were removed from of the -80 °C freezer, thawed on ice, and diluted to the corresponding titer for microinjection to the brain.

### Surgery

Before surgery, each mouse was given dexamethasone (3 mg/kg, i.m.; DexaJect, 002459, Henry Schein Animal Health) and carprofen (5 mg/kg, s.c.; 024751, Henry Schein Animal Health) for anti-inflammatory and analgesic purposes. At the start of surgery, anaesthesia was induced with 2-3% isoflurane and the mouse was affixed in a stereotaxic apparatus (Model 900, David Kopf Instruments). Anaesthesia was maintained with 1-1.5% isoflurane. Body temperature was maintained at 38 °C using a far-infrared warming pad (RT-0515, Kent Scientific). Petrolatum ophthalmic ointment (IS4398, Dechra) was applied to cover the eyes. The scalp was disinfected by wiping with ethanol pads and povidone-iodine. Small burr holes were made above the targeted brain regions using a handheld dental drill (HP4-917, Foredom). AAV was delivered intracranially into the brain by inserting a borosilicate glass capillary and using an injector (Nanoject II Auto-Nanoliter Injector, Drummond Scientific). Injections were performed using 4.6[nL pulses with a 10 s interval between each pulse. To reduce the backflow of the virus, we waited 5-10 min after completing an injection at one site before retracting the pipette to move on to the next site. For the medial frontal cortex, the stereotaxic apparatus was positioned at four sites corresponding to four vertices of a 0.2-mm-wide square centered at the coordinates outlined below. Throughout the procedure, the brain surface was kept moist with artificial cerebrospinal fluid (aCSF; 135 mM NaCl, 5 mM HEPES, 5[mM KCl, 1.8 mM CaCl_2_, 1 mM MgCl_2_; pH[7.3). After injections, the craniotomies were covered with silicone elastomer (0318, Smooth-On), and the skin was sutured (1265B, Surgical Specialties). At the end of surgery, the animal was given carprofen (5 mg per kg, subcutaneous) immediately and then again on each of the next three days.

To fluorescently label a sparse subset of PT neurons, 110.4 nL of AAVretro-hSyn-Cre-WPRE-hGH (1:100 diluted in PBS (P4417, Sigma-Aldrich)) was injected into the right pons (anteroposterior (AP), -3.4[ mm; mediolateral (ML), -0.7 [mm; dorsoventral (DV), -5.2 mm; relative to bregma) and 147.2[nL of AAV1-CAG-Flex-mRuby2-GSG-P2A-GCaMP6f-WPRE-pA (1:10 diluted in PBS) was injected into the ACAd and medial MOs subregion of right medial frontal cortex (AP, 1.5 mm; ML, -0.4 mm; DV, -1.0 mm; relative to bregma) of each C57BL/6J, *Htr2a^f/f^*, or *Htr1a^f/f^*mouse. This procedure yields sparse fluorescent labeling of PT neurons in the medial frontal cortex. We note that, in *Htr2a^f/f^* and *Htr1a^f/f^*mice, the low-titer AAVretro-hSyn-Cre would induce knockout in additional pons-projecting cells that are not fluorescently labeled. After 2-3 weeks, the mouse underwent a second procedure, with the same pre- and post-operative care, to implant a glass window for imaging. An incision was made to remove skin above the skull, and the skull was cleaned to remove connective tissues. A dental drill was used to make an approximately 3-mm-diameter circular craniotomy above the previously targeted location at the medial frontal cortex. aCSF was used to bathe the exposed dura in the craniotomy. A two-layer glass window was made by bonding two round coverslips (3 mm diameter, 0.15[mm thickness; 640720, Warner Instruments) with ultraviolet light-curing optical adhesive (NOA 61, Norland Products) using an ultraviolet illuminator (2182210, Loctite). The glass window was placed over the craniotomy and, while maintaining a slight pressure, super glue adhesive (Henkel Loctite 454) was carefully used to secure the window to the surrounding skull. A stainless steel headplate (eMachineShop; design available at https://github.com/Kwan-Lab/behavioural-rigs) was secured onto the skull and centered on the glass window using a quick adhesive cement system (Metabond, Parkell). The mouse would recover for >10[days after the window implant prior to treadmill habituation and experimental imaging sessions.

### Histology

Histology was performed to determine the accuracy of injection locations and assess transgene expression. For two-photon imaging, after completion of experiments, mice were perfused with PBS, followed by paraformaldehyde solution (PFA, 4% (v/v) in PBS). The brains were extracted and further fixed in 4% PFA at 4 °C for 12-24 h. Subsequently, 40-50-µm-thick coronal sections were obtained using a vibratome (VT1000S, Leica) and mounted onto slides using Vectashield containing DAPI (H-1200-10, Vector Laboratories) and a glass coverslip. Brain sections were then imaged using a wide-field fluorescence microscope (BZ-X810, Keyence).

### Liquid chromatography-tandem mass spectrometry

The timing of brain psilocin distribution and elimination was examined using liquid chromatography-tandem mass spectrometry. Sixty C57BL/6J mice (29 F, 31 M) were treated with a single dose of psilocybin (1 mg/kg, i.p.) and returned to their home cage. At specified time points after drug administration, mice were euthanized and their brain extracted. Eight groups were collected for the time points of 5 minutes (n = 7; 3 F, 4 M), 10 minutes (n = 7; 4 F, 3 M), 20 minutes (n = 6; 3 F, 3 M), 30 minutes (n = 8; 4 F, 4 M), 1 hour (n = 6; 3 F, 3 M), 2 hours (n = 6; 3 F, 3 M), 4 hours (n = 8; 3 F, 5 M), and 24 hours (n = 6; 3 F, 3 M). For controls, 6 mice (3 F, 3 M) did not receive an injection. The brain tissue was immediately homogenized with ascorbic acid (25 mM; PHR1008, Millipore-Sigma) to prevent oxidation [61]. Samples were then flash-frozen and stored at -80*°*C until further processing. The presence of psilocin in brain tissue was determined using liquid chromatography-tandem mass spectrometry (Nexera Prominence, Shimadzu; ZenoTOF 7600, SCIEX) in the Proteomics and Metabolomics Facility at the Cornell Institute of Biotechnology. The peak area detected in the brain tissue was normalized using an external standard for psilocin (#9003135, Cayman Chemical). The timecourse of psilocin brain concentration following psilocybin administration was then fit using a one-compartment pharmacokinetic model:

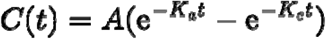

Where *C* is brain psilocin concentration, *t* is time, *A* is a constant, *K_a_* is the absorption rate, *K_e_* is the elimination rate. From the fitted curve, we determined the peak concentration *C_max_*, time to peak *t_max_*, and elimination half-life, *t_1/2_*.

### Head-twitch responses

Head-twitch response data were from our previous study [47]. Briefly, head movements were measured in adult C57BL/6J mice (n = 6; 3 M, 3 F) for 120 minutes after psilocybin treatment (1 mg/kg, i.p) using a magnetic ear tag detection system. We determined the rate of head twitch response by calculating the running average using a 3-minute sliding window.

### Two-photon imaging

Two-photon imaging experiments were performed using a Movable Objective Microscope (MOM, Sutter Instrument) equipped with a resonant-galvo scanner (Rapid Multi Region Scanner, Vidrio Technologies) and a water-immersion ×25 objective (CFI APO LWD, ×25/1.10[NA, Nikon). ScanImage 2020 software [62] was used to control the microscope for image acquisition. To visualize the PT neurons co-expressing mRuby2 and GCaMP6f, two femtosecond-pulsed lasers were aligned to converge into the excitation path: a fixed-wavelength laser for excitation at 1064 nm (ALCOR-1064-2W with XSight Module, SPARK Lasers), and a fixed-wavelength laser for excitation at 920 nm (Axon 920-2 TPC, Coherent).

Structural imaging of mRuby2 was acquired using only the 1064 nm excitation path, whereas calcium imaging of GCaMP6f was done using only the 920 nm excitation path. For all imaging, emitted fluorescence signals were collected through a 580-680 nm bandpass filter for mRuby2 and a 475-550 nm bandpass filter for GCaMP6f. For all experiments, the laser power measured at the objective was ≤120[mW and varied depending on the imaging depth. For longitudinal imaging of the same field of view across days, the laser power was kept the same at each imaging session.

We imaged GCaMP6f signals from the same dendritic branches longitudinally for three days before, during, and after treatment with psilocybin (1 mg/kg, i.p.) or saline (10 mL/kg, i.p.). To target the ACAd and medial MOs subregion of the medial prefrontal cortex, all imaging fields of view were within 500 µm of the midline as determined by first visualizing the sagittal sinus in bright-field imaging. To target apical tuft dendrites of single neurons, we first imaged 0-200 µm below the pial surface to identify candidate dendritic branches. Tuft dendrites were then followed down to their corresponding apical trunk and soma to confirm PT neuron morphology and branching hierarchy. The sparse viral labeling strategy allowed definitive assignment of each apical dendritic branch to the neuron. An imaging field of view was then selected between 20-150 µm below the pial surface, to capture apical tuft dendrites that were at least three branch points distal to the apical dendritic trunk. In most cases, all apical dendrites in a field of view originated from one neuron. On day -1, the day before treatment, a baseline calcium imaging session of 15-20 minutes was acquired. On day 0, the same field of view was imaged again for 30 minutes pre-treatment (“Pre”). Imaging was then paused to inject psilocybin or saline. Within 1 minute of the injection, imaging was resumed for 90 minutes (“Post”). On day 1, the day after treatment, the field of view was imaged again for 60-90 minutes (“24hr”).

All time-lapse calcium imaging data were acquired with 30 Hz bidirectional scanning at 512 ×[512 pixels with resolutions of 0.497-1.000 µm per pixel. A subset of mice underwent both psilocybin and saline treatment, with at least 10 days between treatments. Altogether, 21 C57BL/6J mice (11 F, 10 M) were treated with psilocybin (84 dendritic branches, from 13 cells) and saline (90 branches, from 15 cells); 9 *Htr2a^f/f^* mice (7 F, 2 M) were treated with psilocybin (90 branches, from 11 cells) and saline (56 branches, from 7 cells); and 6 *Htr1a^f/f^* mice (3 F, 3 M) were treated with psilocybin (45 branches, from 5 cells) and saline (26 branches, from 4 cells). In rare cases, microscope fault (e.g., objective lost water immersion, power outage) caused early termination of an imaging session with at least half the session completed. Data for these interrupted sessions were retained with dropped time points treated as missing (i.e., one C57BL/6J mouse/cell at pre, one C57BL/6J mouse/cell at post). One C57BL/6J mouse/cell was unable to be imaged at the 24hr hour timepoint, so its data for that full session was treated as missing.

For structural imaging of dendrites on day -1 and day 1, the mouse was lightly anaesthetized with 0.5-1% isoflurane through a nose cone after completion of calcium imaging. Each structural imaging session lasted 20-30 min. For each field of view, a 50- to 100-µm-thick *z* stack centered on the field of view used for calcium imaging was collected with 1-µm steps using 15[Hz bidirectional scanning at 1,024[×[1,024 pixels and resolutions of 0.333-0.427 µm per pixel.

These structural images allowed for the determination of each dendritic branch with a relative position in its apical tuft based on landmarks (i.e., branch points) not seen in the planar calcium imaging fields of view. At the end of the imaging session on day 1, for coarse reconstruction of the entire PT neuron, a *z*-stack was acquired between 0 and 800 µm below the dura with 1-5[µm steps.

### Linear treadmill

All two-photon imaging experiments were conducted with the head-fixed mice free to run on a low-friction rodent-driven treadmill (Janelia 2017-049 [63]). Each mouse habituated to the treadmill for 3-5 days, with increasing duration each day, prior to imaging sessions. Habituation took place in an enclosure built to mimic the two-photon imaging rig. The number of sessions varied based on each mouse learning to run forward on the treadmill. Mice that did not run forward spontaneously after habituation were not used in this experiment. Before advancing to the experiment, each mouse had at least one habituation session longer than the longest duration of an imaging session.

Treadmill speed was acquired by a single rotary encoder at 1 kHz and synchronized to the imaging frame clock using a two-channel digitizer (MiniDigi 1B, Molecular Devices). The average treadmill speed was calculated over each imaging frame to generate a speed timecourse at the imaging frequency. This timecourse was filtered for a shifting baseline due to belt tripping inconsistent with running. Observations were qualitatively unchanged without these data processing steps. Locomotor timecourses were then used to distinguish epochs of rest (speed = 0 cm/s) and run (speed > 0 cm/s). The distance traveled was calculated as the integral of the speed timecourse. To test for drug-evoked changes in locomotion, the average pre-treatment distance traveled per 10 minutes was subtracted from the distance traveled in each 10-minute time bin pre- and post-treatment.

### Analysis of the imaging data

For dendritic calcium imaging, the image files from each experiment were processed with PatchWarp [64] in MATLAB to correct for non-rigid translational motion within each experimental session (i.e., day -1, pre, post, 24hr). Motion correction was performed based on the static fluorescence signals in the mRuby2 channel, then applied to the data from the GCaMP6f channel. For longitudinal alignment across sessions, data from all sessions were aligned to a template generated by averaging the motion-corrected baseline session data (day -1). Inter-session alignment was confirmed for all sessions by manual inspection. As an overview, the additional processing steps included: (1) manually tracing regions of interest (ROIs) corresponding to dendritic branches using an in-house graphical user interface in MATLAB; (2) extracting the average fluorescence trace from each ROI and process similar to previous work [13, 65] to exclude background neuropil signal, and convert to fractional change in fluorescence (Δ*F*/*F*(t)); (3) deconvolving the fluorescence trace into discrete calcium event probabilities. These processing steps are detailed below. To trace dendritic branch ROIs, the field of view from baseline session was used, while checking the corresponding structural z-stack images to identify dendritic segments originated from the same PT neuron. First, a mock ROI would be traced along a dendritic shaft segment using a lasso drawing tool. The mock ROI was then overlayed on the average image of each calcium imaging session to check for longitudinal agreement. The ROI was then redrawn or edited until it captured the greatest extent of shaft that was consistently in the focal plane across sessions. In this way, a single ROI mask was used to extract calcium signals for each experimental calcium imaging session. For each ROI, the pixel-wise average fluorescence was calculated at each data frame to generate a timecourse *F*_ROI_(t) per imaging session. All ROI tracing was performed by an experimenter blinded to the treatment group.

Each ROI was then processed to reduce the contribution from background neuropil. Each planar branch ROI was dilated by *r* in all directions, and an ROI-specific neuropil mask was created by subtracting a branch ROI dilated by *r* from a second branch ROI dilated by 2*r* (where *r* = 2.5 µm). Neuropil masks excluded pixels belonging to any other dendritic branch ROI. Finally, the remaining pixels in the neuropil mask were averaged per data frame to generate *F*_neuropil_(t). Each ROI had the fluorescence from its neuropil mask subtracted as follows:

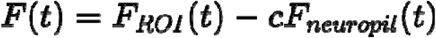

where the neuropil correction factor, *c*, was set to 0.4. Next, the fractional change in fluorescence Δ*F*/*F*(t) was calculated for each ROI by normalizing *F*(t) to its baseline, *F*_0_(t), estimated as the 10th percentile within a 2-minute sliding window:

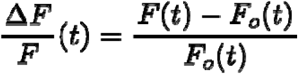

Calcium events were detected from the Δ*F*/*F*(t) of each ROI using a generalized spike inference toolbox, CASCADE [46]. Briefly, CASCADE includes deep network models that are trained to detect absolute neuronal spike rates based on diverse ground truth experiments of calcium indicators with simultaneous electrophysiological recordings from neurons. CASCADE has been shown to require minimal parameter optimization while achieving high performance. For each Δ*F*/*F*(t) timecourse, a spike probability timecourse was generated using CASCADE’s pretrained network model, “Global_EXC_30Hz_smoothing25ms_causalkernel”. Spike probability timecourses represent the expected number of spikes occurring in each imaging frame. Each spike probability timecourse was visually inspected against its originating Δ*F*/*F*(t) timecourse to confirm precision.

For each imaging session, an ROI’s calcium event rate was computed by summing the spike probability timecourse and dividing by the duration of the imaging session. The change in calcium event rate was computed for each ROI by calculating post-treatment rate minus pre-injection rate divided by the pre-treatment rate. To calculate event rates separately for rest and run, the spike probability timecourse was summed within the image frames labeled as rest or run, then divided by the total time spent in rest or run (e.g., sum of spike probability during rest frames divided by the total time labeled as rest). To determine dendritic calcium events as a function of time, spike probability timecourse was summed in one-minute bins. In this case, the change in calcium event rate as a function of time was computed for each ROI by calculating the rate in the current one-minute bin minus pre-injection rate divided by the pre-treatment rate (e.g., tenth minute post-treatment rate minus full pre-treatment rate divided by the full pre-treatment rate).

For structural imaging of dendrites, a subset of the imaged dendritic branches (n = 113 branches from 21 neurons from 21 C57BL/6J mice, 10 M, 11 F) showed sufficient mRuby2 fluorescence to reliably quantify dendritic spines across time. For these samples, an experimenter blinded to treatment condition quantified the number of dendritic spines detected on day -1 and day 1. To match the structural imaging to calcium imaging, the experimenter used the dendritic branch ROI mask from the calcium imaging as a reference for the proximal and distal limits of the dendritic segment within which to measure spines. 105 of the 113 dendritic branches with tracked structure had complete a Ca^2+^ imaging dataset for comparison across modalities. To measure spine density, dendritic spines were counted using standardized criteria [66] as done previously by our lab [11, 13]. Briefly, a dendritic spine was counted when the protrusion extended for >0.4 μm from the dendritic shaft. The line segment tool in ImageJ was used to measure the distances. The spine density was calculated as the number of spines counted per unit length of dendritic branch segment. For each dendritic segment, the change in spine density was computed by calculating the spine density on day 1 minus the spine density on day -1 divided by the spine density on day -1.

### Statistics

All statistical tests were computed with R and MATLAB. Sample sizes were based on pilot and existing relevant studies and were not statistically predetermined. All tests were two-sided, and results are displayed as the mean[± bootstrapped 95% confidence intervals of the mean (resampled with replacement 1000 times). Linear mixed-effects models were used to estimate the effects of treatment, time, genotype, and locomotion on drug-evoked changes in dendritic calcium events using the lme4 package in R. *Post hoc* analysis was performed with *t*-tests using the Bonferroni method to correct for multiple comparisons. Mixed-effects models were preferred to traditional repeated measures ANOVAs due to their advantages in datasets where repeat measures are influenced by within-subject nesting (for example, multiple branches per cell per mouse) and instances of missing data. Pearson’s correlations were used to test the relation between psilocybin’s pharmacokinetics, head twitch responses, and drug-evoked changes in dendritic calcium events. Pearson’s correlations were also used to quantify the relationship between changes in dendritic spine density and changes in dendritic Ca^2+^ events. The procedures for statistical testing are detailed below, with all models and *post hoc* testing outputs reported in **Table S1**.

## Data and code availability

All data tables and code for analyses presented in this paper will be deposited in Github and made publicly available at the date of publication. Any additional information required to reanalyze the data reported in this paper is available from the lead contact upon request.

## ACKNOWLEDGEMENTS

Psilocybin was provided by Usona Institute’s Investigational Drug & Material Supply Program, which is supported by Alexander Sherwood, Robert Kargbo, and Kristi Kaylo in Madison, WI. We thank the Genomics Facility (RRID:SCR_021743) of the Biotechnology Resource Center of Cornell Institute of Biotechnology for their help with LC-MS/MS experiments. We thank Hail Kim and Takeshi Sakurai for providing *Htr2a^f/f^*and *Htr1a^f/f^* mouse lines, respectively. The authors acknowledge the support of National Institute of Health grants R01MH128217 (A.C.K.), R01MH137047 (A.C.K.); the One Mind–COMPASS Rising Star Award (A.C.K.); NIH training grant T32GM007205 (N.K.S.); National institute of Health fellowships F30MH129085 (N.K.S.) and F31EY038625 (C.A.K.); and NIH instrumentation grant S10OD032251 (Cornell Biotechnology Resource Center Imaging Facility).

## CONTRIBUTIONS

N.K.S. and A.C.K. planned the study. N.K.S. conducted and analyzed two-photon imaging experiments. L.-X.S. analyzed the structural imaging data. C.A.K. and A.D.G. conducted and analyzed the HPLC-MS experiments. Q.J. assisted with histology. N.K.S. and A.C.K. drafted the manuscript. All authors reviewed the manuscript before submission.

## SUPPLEMENTAL INFORMATION

**Table S1.** Comprehensive statistical details for all data presented in the article.

**Document S1.** Supplementary figures and legends (Figures S1–S4).

