## Supplementary figures for "Psychedelic drug action at dendrites is gated by behavioral state and serotonin receptors"

**This PDF file includes:**

Supplementary figures and legends (Figures S1 to S4)

**Other supporting materials for this manuscript include:**

Table S1 (.xlsx)

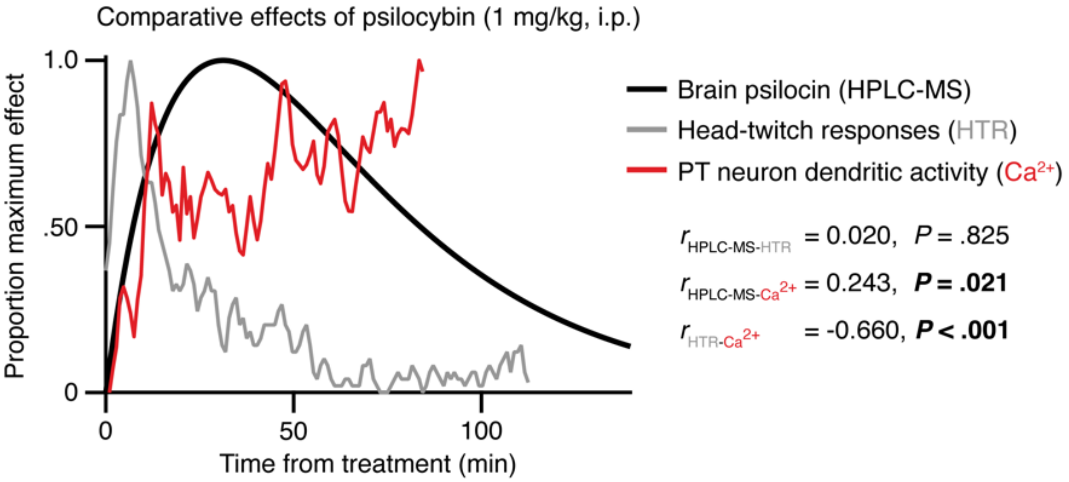


**Figure S1. Psilocybin-evoked dendritic calcium activity correlates with brain psilocin concentration, but not with head-twitch response.**

We compared the acute effect of psilocybin treatment (1 mg/kg, i.p.) on brain psilocin concentration (i.e., measured by HPLC-MS), head-twitch response (HTR), and the drug-evoked increase in calcium events at the dendrites of PT pyramidal neurons. Data for HTR came from our lab's previous study (Aboharb et al., Nat. Commun., 2025).

Psilocybin increases dendritic calcium events in frontal cortical PT neurons, over a time course that correlated with the brain psilocin concentration ($r = 0.24$, $P = 0.021$). By contrast, HTR occurred much earlier, peaking 6-8 minutes after treatment. Note that although there was a significant correlation coefficient between HTR and dendritic calcium change, the correlation coefficient is negative ($r = -0.66$).

For HPLC-MS, n = 60 mice (31 F, 29 M), 6-8 mice per time point (5, 10, 15, 30, 60, and 120 min after treatment; later time points not used in this analysis). Data were fit with a one-compartment pharmacokinetics model. For HTR, n = 6 (3 F, 3 M), 0-120 min after treatment. For dendritic calcium activity, n = 84 dendritic branches from 13 PT neurons, 1-90 min after treatment. Full statistical reporting is provided in **Table S1**.

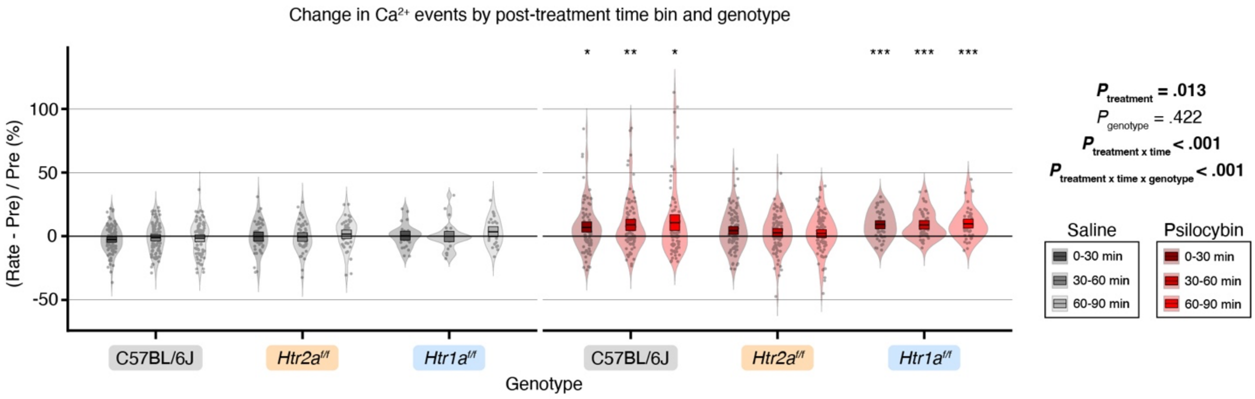


**Figure S2. 5-HT$_{2A}$ and 5-HT$_{1A}$ receptors exert diverging effects on the psilocybin-evoked increase in dendritic dynamics.**
Fractional change in calcium events detected in PT neuron dendritic branches in 30-minute time bins after treatment with psilocybin (red) or saline (black), across genotypes. Branch-wise distributions are shown in violins, with the bootstrapped 95% confidence interval of the mean inlaid as a box. Statistics to the right summarize select fixed effects from a mixed model on the minute-wise imaging data. Asterisks denote significance on one-sample *t*-tests (mean vs. 0% change) after Bonferroni correction for multiple comparisons across time and genotype: * $P < 0.05$, **, $P < 0.01$, *** $P < 0.001$.

For C57BL/6J, n = 84 and 90 dendritic branches from 13 and 15 PT neurons for psilocybin and saline, respectively. For *Htr2a*$^{f/f}$, n = 87 and 45 dendritic branches from 11 and 7 PT neurons for psilocybin and saline, respectively. For *Htr1a*$^{f/f}$, n = 45 and 26 dendritic branches from 5 and 4 PT neurons for psilocybin and saline, respectively. Full statistical reporting is provided in **Table S1**.

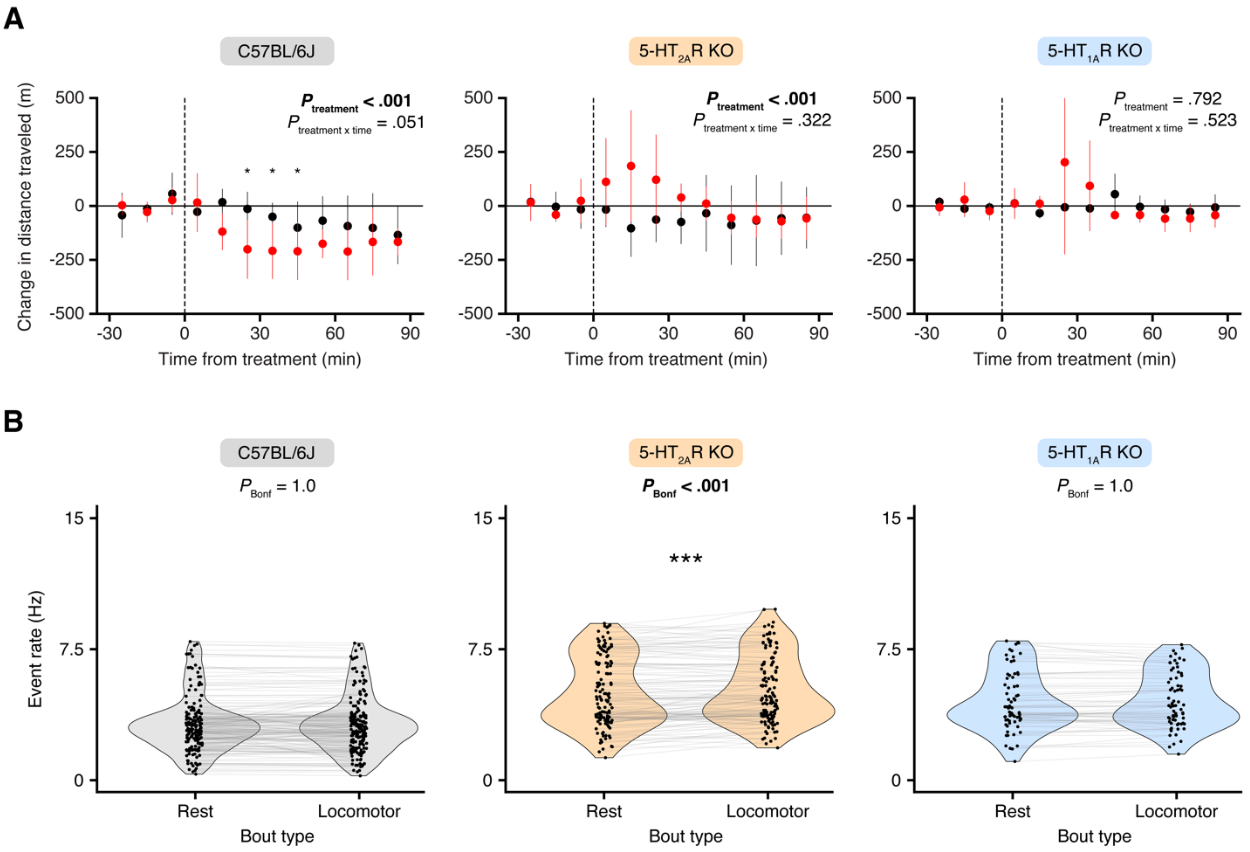


**Figure S3. Locomotor activity and dendritic calcium events across genotypes.**
**(A)** Changes in the distance traveled in 10-minute time bins on the treatment day (Day 0) for each mouse strain. Error bars, the bootstrapped 95% confidence interval of the mean. *P* values from post-hoc *t*-tests were adjusted using Bonferroni correction for multiple comparisons across time. *, $P < 0.05$.
**(B)** Prior to treatment, the rate of dendritic calcium events during rest and run epochs. Circle, individual dendrite. *P* values from post-hoc paired-samples *t*-tests were adjusted using Bonferroni correction for multiple comparisons across genotype. ***, $P < 0.001$.

For (A), n = 21 C57BL/6J mice (11 F, 10 M); n = 9 *Htr2a*$^{f/f}$ mice (7 F, 2 M); n = 6 *Htr1a*$^{f/f}$ mice (3 F, 3 M). For (B), for C57BL/6J, n = 174 dendritic branches from 28 PT neurons; for *Htr2a*$^{f/f}$, n = 132 dendritic branches from 18 PT neurons; for *Htr1a*$^{f/f}$, n = 71 dendritic branches from 9 PT neurons. Full statistical analyses are provided in **Table S1**.

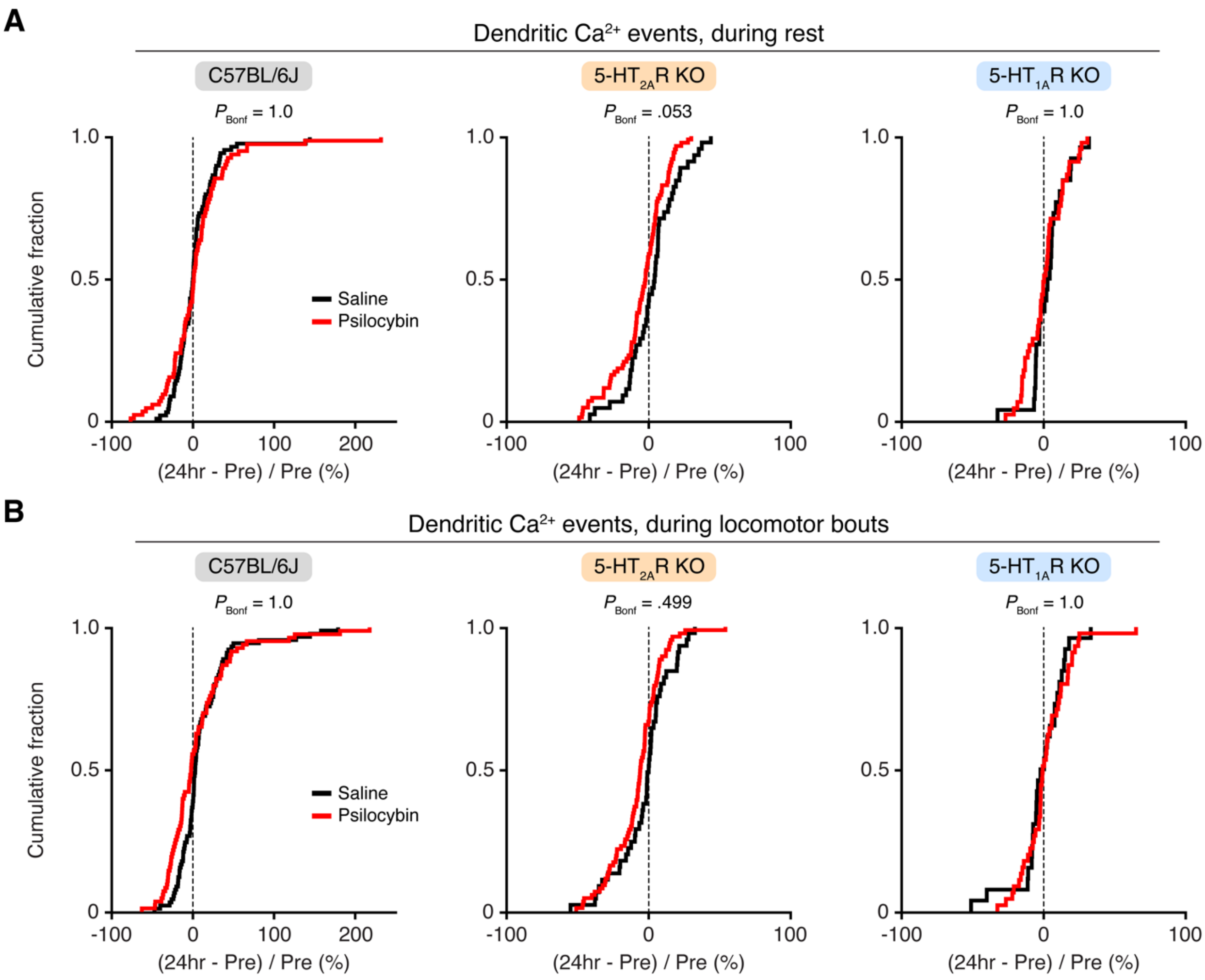


**Figure S4. No effect on the dendritic calcium responses at 24 hours after psilocybin administration, regardless of the genotype.**
**(A)** Cumulative distribution plot of the fractional change in the rate of calcium events detected in PT dendritic branches, restricting analyses to rest epochs, after psilocybin (red) or saline (black) the day after treatment ("24hr") relative to pre-treatment baseline ("Pre"), for C57BL/6J mice (left), *Htr2a$^{f/f}$* mice with 5-$HT_{2A}$ receptor knockout (middle), and *Htr1a$^{f/f}$* mice with 5-$HT_{1A}$ receptor knockout (right). *P* values from post-hoc *t*-tests were adjusted using Bonferroni correction for multiple comparisons.
**(B)** Similar to A, but restricting analyses to run epochs.

For C57BL/6J, n = 83 and 90 dendritic branches from 13 and 15 PT neurons for psilocybin and saline, respectively. For *Htr2a$^{f/f}$*, n = 87 and 45 dendritic branches from 11 and 7 PT neurons for psilocybin and saline, respectively. For *Htr1a$^{f/f}$*, n = 45 and 26 dendritic branches from 5 and 4 PT neurons for psilocybin and saline, respectively. Full statistical analyses are provided in **Table S1**.
